# Shedding light on day-night habitat use of oyster reefs by fish

**DOI:** 10.1101/2025.10.02.680156

**Authors:** Diego L. Gloria, Michael Sievers, Cesar Herrera, Rod M. Connolly

## Abstract

Oyster reefs provide key ecosystem functions, including the provision of habitat for fish. While their importance in supporting fish assemblages is widely accepted, an assessment of intertidal oyster reefs at night remains a knowledge gap. Documenting habitat use at night could reveal new species reliant on these systems and help uncover diel fish movements among structured and unstructured habitats. We hypothesise that habitat use will interact with time and habitat, whereby differences in fish assemblages between oyster reef and unstructured habitats would be greater during the day. In this study, we used infrared-capable remote underwater video stations (RUVS) to sample paired intertidal oyster reefs and unstructured habitat during day and night. Assemblages using these habitats during the day do not match those at night, consistent with diel shifts observed in similar habitats. Consistent with our hypothesis, during the day oyster reefs had higher diversity and richness than unstructured habitat, whereas both habitats had low diversity and richness at night. The high relative abundance of piscivores in oyster reefs during the day suggests reefs create food webs that extend beyond benthic matter and sustain higher trophic levels. Nighttime abundances and diversity in oyster reefs dropped to levels similar to those of unstructured habitats during the day and night, likely a product of temporal niche partitioning and dynamic predation risks. Nighttime studies were important in documenting a wider suite of species and can thus provide a more complete understanding of fish-habitat interactions. Furthermore, this study used novel methods that enabled depth and size estimation from a single camera unit. While still in its infancy, this can serve as a force multiplier in monitoring intertidal habitats and lead towards a cost-effective method for standardising day/night abundance measurements and obtaining fish size estimates.

## Introduction

Oyster reefs provide key ecosystem services including water quality improvement, nitrogen sequestration, coastal defence, and fishery production (Beck et al. 2011; Grabowski et al. 2012; Kroeger 2012). In the exposed intertidal zone, they support biodiversity by providing shade and refuge for invertebrates in an otherwise extreme physical environment (McAfee et al. 2017; Boström-Einarsson et al. 2022). The habitat complexity provided by oyster reefs supports a wide variety of animals ranging from small amphipods to top predators such as sharks, thus creating a diverse food web dependent on this key habitat (Smyth and Roberts 2010; Grabowski et al. 2019; Howie and Bishop 2021). Furthermore, these reefs form part of the ecosystem mosaic that fish can use as both nursery and feeding grounds (Tolley and Volety 2005; Beck et al. 2011; Nagelkerken et al. 2015; zu Ermgassen et al. 2016; Gilby et al. 2018; Martinez-Baena et al. 2023). Indeed, oyster reefs can enhance the biodiversity and abundance of animals both inside and around the reef (De Santiago et al. 2019; Searles et al. 2022).

Despite these key functions, oyster reefs have experienced an 85% decline globally, with many becoming functionally extinct (Beck et al. 2011). The oyster reef ecosystem on Australia’s east coast is classed as critically endangered (Gillies et al. 2020), and Moreton Bay in southeast Queensland has experienced a 100% loss of subtidal oyster reefs and a reduction of 96% in the vertical zonation of oysters (Diggles et al. 2019; Thurstan et al. 2020). One important step in protecting these habitats is to understand the environmental and economic value of oyster reefs to garner support for their conservation and restoration (Barbier 2017). It is estimated that the total annual value of the ecosystem services provided by oyster reefs ranges between US$5500 and US$99000 per hectare (Grabowski et al. 2012). Part of this value is provided by their function in commercial fisheries; previous studies have suggested that habitat provisioning by oyster reefs yields US$20 million annually in an area of 1,045 hectares (Lai et al. 2020) and that a hectare of oyster reef is estimated to have a value of US$4123 annually for commercial fisheries alone (Grabowski et al. 2012). However, these estimates are based on daytime surveys and studies in temperate oyster reefs in the United States. Subtropical and intertidal oyster reefs, including those in Australia, are less studied and an accurate valuation of their contribution to Australian fisheries remains speculative.

Although there is a lack of comprehensive night surveys of oyster reefs, previous studies in seagrass, temperate reefs, and rocky reefs show significant differences between day and night faunal assemblages (Guest et al. 2003; Myers et al. 2016; Larissa et al. 2023), and sampling at night increases the chance of documenting rarer species (Griffiths 2001; Hagan and Able 2007). Light can alter fish behavior and habitat use (Didrikas and Hansson 2008; Bolton et al. 2017) as fish attempt to balance predation risk and foraging needs (Metcalfe et al. 1999).

Predatory species may become more active at night (Shoji et al. 2017) and can exploit structurally complex habitats such as oyster reefs to create ambush opportunities (Michel et al. 2020). This is supported by higher abundances of predators observed at night in estuarine habitats such as seagrasses and muddy sediments (Griffiths 2001; Hagan and Able 2008; Castillo-Rivera et al. 2010). An increase in piscivorous species has also been documented in seagrass beds at night (Shoji et al. 2017), while rocky reefs experience a marked change in the abundances of various feeding guilds between day and night (Larissa et al. 2023).

Furthermore, fish biomass may increase at night in estuaries (Guest et al. 2003; McSpadden et al. 2023). These habitats often coexist with oyster reefs as part of a larger seascape, and understanding coastal ecology requires thorough knowledge of how each of its constituent parts function throughout the entire diel cycle. For example, habitats that serve as nurseries during the day may be feeding grounds at night (Shoji et al. 2017). These studies show the importance of sampling at night to gain a comprehensive assessment of coastal ecosystems, and thus to a better understanding of the benefits of these habitats and the services they provide.

Effective restoration requires the integration and knowledge of key animals that use the habitat (Reeves et al., 2020; Howie & Bishop, 2021; Sievers et al., 2022), and funding and support from stakeholders often relies on a tangible value for ecosystem services. Accurate estimates of fish use across day and night may thus improve the support and management of oyster reefs (Barbier 2017). Furthermore, having suitable reference data to help form the basis of oyster reef conservation is important for guiding restoration and measuring success (Gann et al. 2019; Fitzsimons et al. 2020). Current data solely based on daytime surveys may underestimate or overestimate the value of oyster reefs to fisheries and as habitats and may mean that our reference information for oyster reefs is incomplete. Using non-invasive methods at night allows for the observation of natural fish behavior (Mallet and Pelletier 2014) that has not yet been documented for oyster reefs. Surveying at night may also add to the known list of species that rely on oyster reefs, which can create a more complete baseline for future work while also incentivising restoration. By using infrared-light sensitive cameras, a more accurate image of oyster reefs can be painted to further inform both future nighttime studies as a whole and oyster reef ecology in particular.

With previous studies in other coastal habitats showing abundance changes in functional groups and shifts in habitat functions from day to night, we hypothesised that the abundance and diversity of fishes would interact with habitat type (oyster reef vs. unstructured) and time (day vs night). Unstructured habitats served as controls, allowing us to distinguish general patterns of habitat use from the unique ecological contributions of oyster reefs. Specifically, we anticipated that the difference between the two habitats would be greater at daytime when a wider range of fish are more likely to be active and using the structured reefs, while at night the difference would be reduced, potentially due to increased risk of predation (Shoji et al. 2017; Michel et al. 2020).

## Methods

### Study Areas

We surveyed fish assemblages in 13 intertidal oyster reefs in three estuaries spanning Pumicestone Passage in northern Moreton Bay (−27.042250, 153.115930) to Tallebudgera Creek (−28.108321, −153.450358) in the south (Figure 1). The oyster reefs were primarily composed of the Sydney rock oyster (*Saccostrea glomerata*) with a maximum depth of 2.2 m at high tide and were fully exposed at low tide. Oysters were growing on either rock or biogenic oyster remains. To account for possible variations in water depth and visibility, all surveys were conducted within 2 hours prior to the high tide (Gilby et al. 2021). This also maximised the area of the intertidal reef available for fish to use (Mosman et al. 2023).

**Figure 1.**
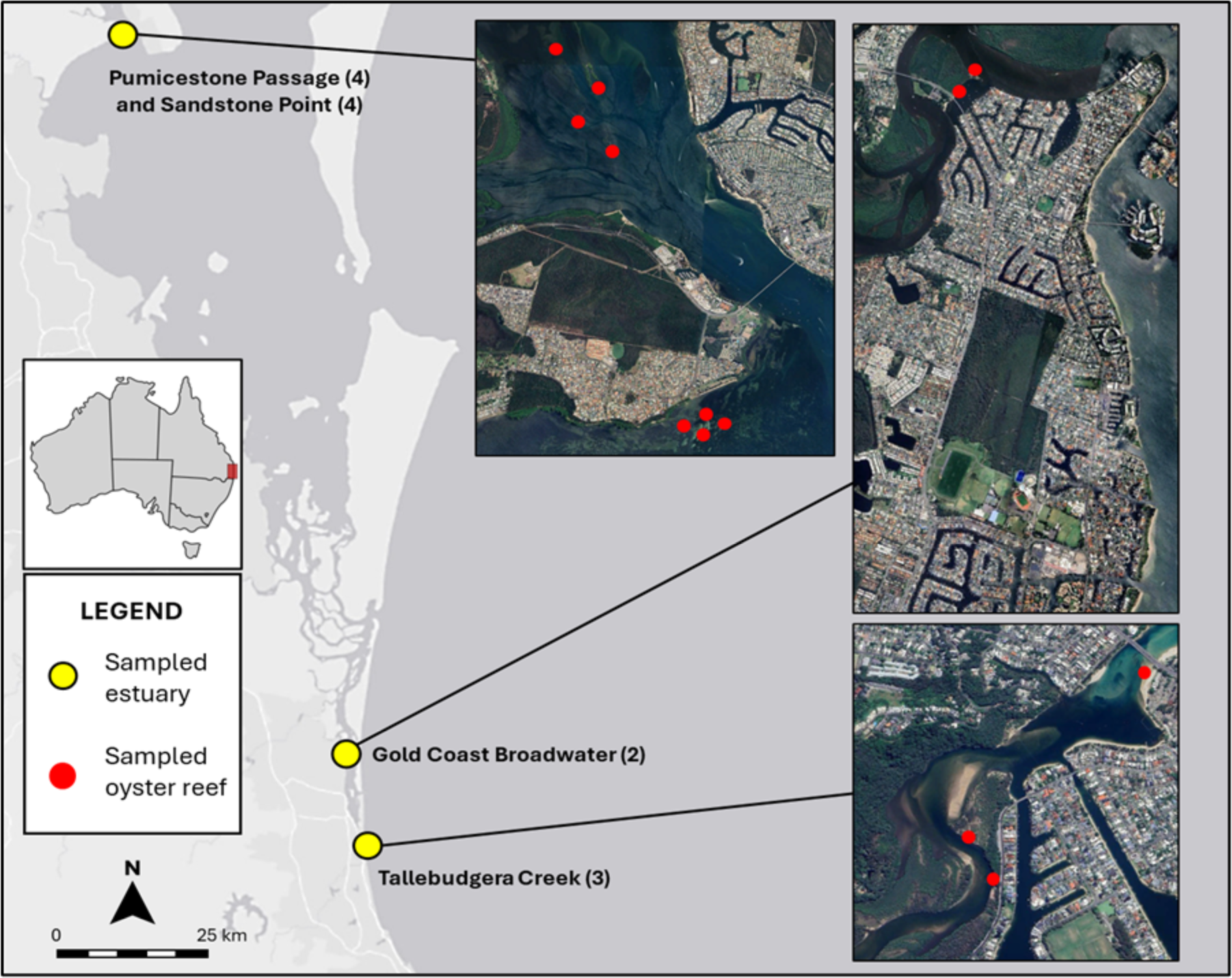
Location of 13 sampled oyster reefs in southeast Queensland (see Table S1 for site coordinates). The number of oyster reef sites sampled in each estuary are noted in parentheses. Each oyster reef was paired with a nearby unstructured control site.

### Sampling Regimes

Each site (N = 13) consisted of a paired oyster reef habitat and an adjacent unstructured habitat to help account for effects caused by environmental variables among sites. The unstructured habitats were used as a control, similar to previous studies on fish using oyster reefs (Connolly et al. 2024; Martin et al. 2024). Oyster reef and unstructured habitats were separated by at least 50 m and were sampled simultaneously. This is greater than the spacing normally used in oyster reef studies (Davenport et al. 2022; Martínez-Baena et al. 2022) and in edge effect studies in other coastal habitats (Mahoney et al. 2018; Carroll et al. 2019; Muething et al. 2020), thus minimising the chances of the unstructured video sample being influenced by the presence of a nearby oyster reef.

Study sites within the same estuary were at least 150 m apart, and sampled simultaneously (i.e., two paired sites per night) making it unlikely that the same fish was sampled twice at two sites. Sites were sampled in both day and night within the same 24-hour window to reduce any confounding effects that may be caused by changing environmental conditions. We defined night as the period at which the sun was 18 degrees or lower from the horizon with no sunlight. This paired design meant that each site was represented by four video recordings (Figure 2).

**Figure 2.**
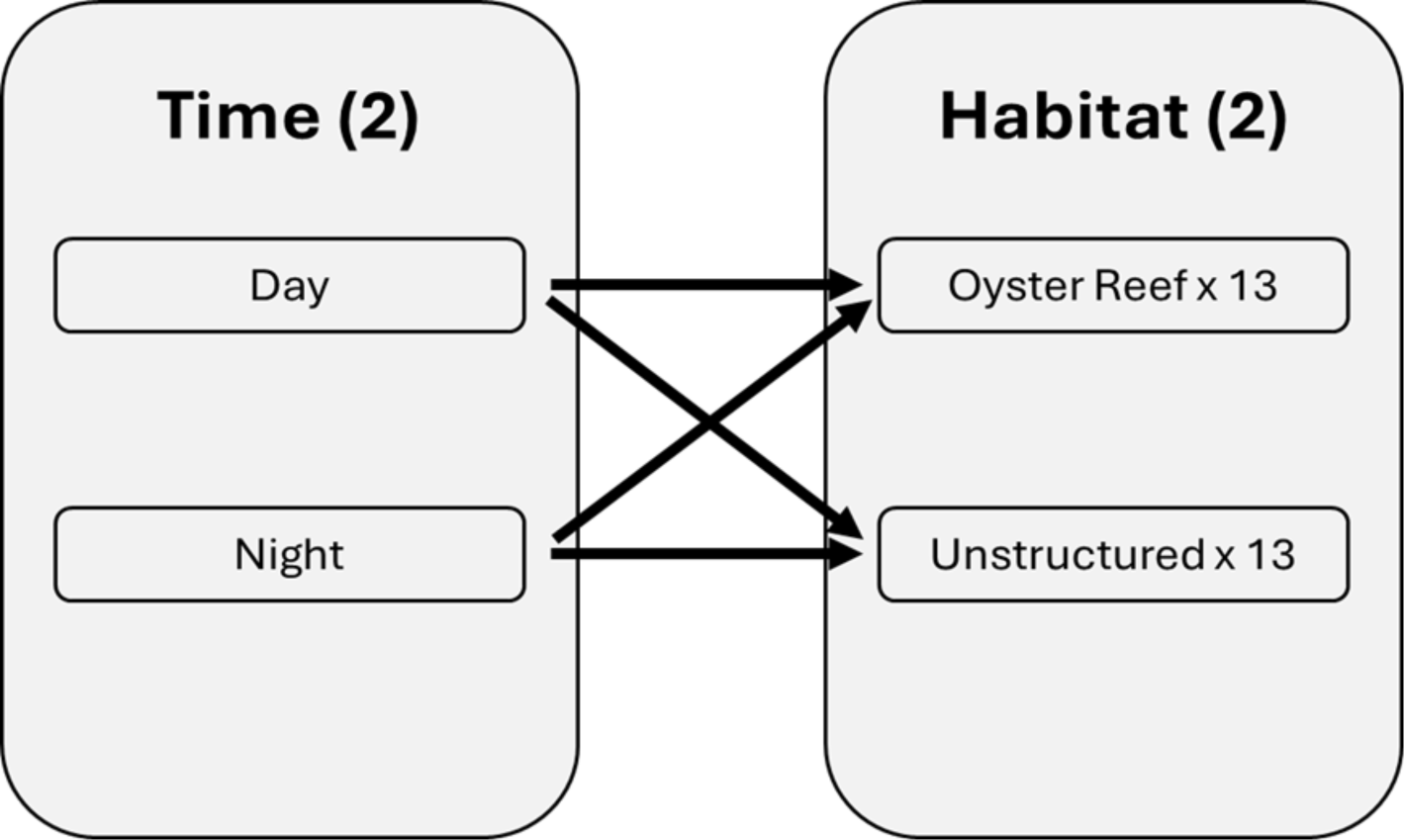
Experimental design showing the paired structure of video surveys. Each site replicate consists of a pair of oyster and unstructured habitats, which were sampled in both day and night. In total, 13 replicates were surveyed.

### Remote Underwater Video Stations

Unbaited remote underwater video stations (RUVS) were used to record fish. This method is non-invasive and allows for natural occurrences and behaviour of fish to be observed (Mallet and Pelletier 2014) without the biases associated with the use of bait (Hardinge et al. 2013; Myers et al. 2016). Each video survey lasted for 60 minutes to adequately represent fish assemblages in the habitat (Erickson et al. 2023). The cameras used were also infrared capable (i.e infrared blocking lens filter removed), as infrared lights were used to illuminate the field of view to allow for video recording at night (Bosiger and McCormick 2014). Most marine animals are insensitive and likely do not detect this spectrum of light (Weiss et al. 2006; Fitzpatrick et al. 2013), compared to other more visible wavelengths that may bias behaviour (Harvey et al. 2012). The infrared lights (turned off) were left on the RUVS during daytime deployments as a procedural control.

RUVS consisted of a single SJ Camera 4000 fixed to a platform with two forward facing infrared lights fixed to either side (Figure 3). A 1.5-metre secchi pole was fixed such that it protruded from underneath the camera to determine the visibility for each deployment (Baker et al. 2022). This provided a cutoff distance to use when counting fish in daytime deployments to account for the lowered visibility at night. Furthermore, the marks on the secchi pole at known distances were used to calibrate depth images created by an AI model for sizing estimation.

**Figure 3.**
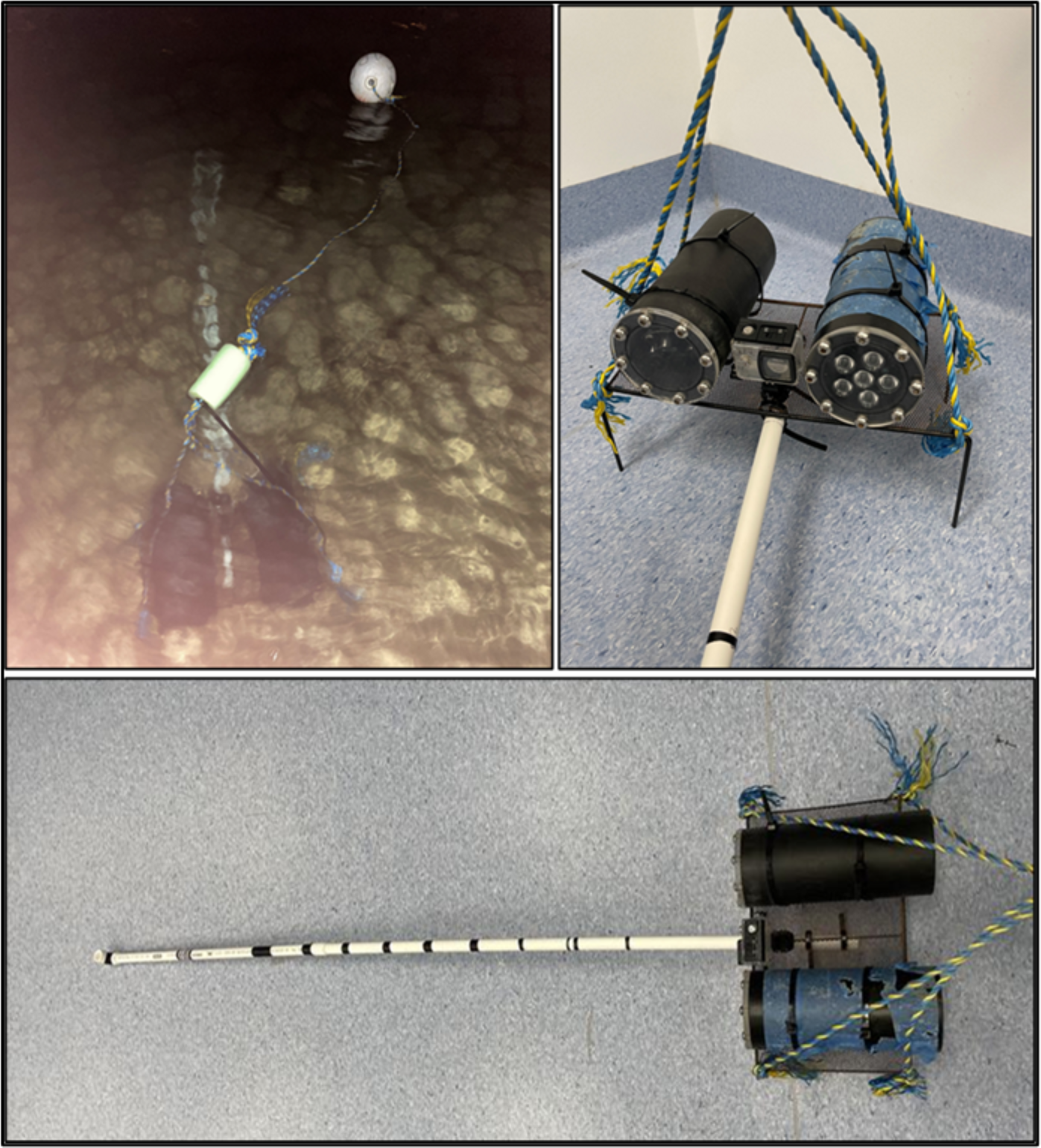
Images of the RUVS unit with infrared lights and a secchi pole outside of and in the water.

RUVS in oyster reef habitats were placed so that the field of view was angled obliquely at the edge of the habitat, thus preventing the habitat structure from obscuring the camera’s field of view. At the start of each deployment, a separate calibration stick with a known length of 30 cm was waved in front of the camera to aid in ground-truthing images for monocular fish sizing.

### Measuring Abundance Using Monocular Depth Estimation and Sizing

To measure fish abundance, the MaxN metric was used. This is defined as the maximum number of individuals of a species visible in one frame of the video and thus serves as a conservative measure of abundance by avoiding double-counts (Myers et al. 2016).

To account for differences in visibility between day and night deployments, a monocular depth estimation artificial intelligence (AI) model was used (Yang et al. 2024). This AI model predicts the distance of objects in a scene using a single image from a single camera, as distinct from the more typical stereoscopic system which requires images from two cameras. The model uses visual cues such as shadows, textures, blur, spatial patterns and colors for assessing the relative geometry of a scene (Bhat et al. 2023). From this information, it is then possible to calculate the distance from objects (i.e., fish) to the camera. To account for the challenging environment posed by underwater scenes with low visibility (e.g. night or high turbidity), each depth map was calibrated by annotating marks of known distances on a reference Secchi pole observable in the image. These annotations were used to build a regression model, which allowed us to estimate the distance to the camera in millimetres (C. Herrera, personal communication, 28 May 2025). Later, fish size was calculated using a simplified pinhole camera model which related the fish’s relative size on the image, the camera focal length, and the estimated distance from the fish to the camera (Hartley and Zisserman 2004). This approach assumed a straight-line projection from the fish to the camera sensor, allowing for the scaling of the observed size to its metric dimensions.

To enable fair comparisons between day and night deployments, fish counts were standardised to avoid overestimating abundances during the day, when higher visibility could result in detecting fish farther away. Visibility during night deployments at each site was measured using the Secchi pole, and this distance was then used as a cutoff distance for counting fish during the corresponding daytime deployment.

The novelty of this technology combined with variable water conditions meant that the RUVS design and deployment methods had to be fine-tuned throughout this study, resulting in reliable depth maps for only 13 deployments across 5 sites. We compared the MaxN obtained by a human observer to the monocular depth results (i.e, MaxN estimates within the distance cutoff value used by the human) in these 13 deployments with reliable estimates using a paired T-test. There was no significant difference (p>0.05; R^2^ = 0.46), so we decided to rely solely on the human MaxN estimates and include the maximum possible 13 sites.

### Data Extraction and Analysis

We analysed 52 hours of video from the 13 sites, each represented by two habitats (oyster reef / unstructured) and two times (day / night). For all models, site was treated as a random factor to account for variability among sites that may have been caused by environmental factors, whilst habitat and time were fitted as interacting fixed effects.

To test for differences in species assemblages, a PERMANOVA using the *adonis2* function with Jaccard distance was used. Rare species that occurred in only 1 deployment, and deployments with no fish observations, were removed to improve dispersion and model fit. Species richness and Shannon’s diversity were computed, and a linear mixed model and analysis of variance (ANOVA) was used to elucidate differences between treatments (Habitat: 2 levels; Time: 2 levels). We used Q-Q and residual plots to confirm whether the models met appropriate statistical assumptions. Where needed, transformations were performed. Post-hoc tests for these metrics were conducted using the estimated marginal means package (*emmeans*).

Each species was then tested individually for differences in abundance. Due to the statistical properties of MaxN and the high proportion of zeros in our data, an Aligned Rank Transform (ART) ANOVA (Kay et al. 2025) with the same model structure as above was used for species with abundances greater than 10, while a Fisher’s exact test was used on the remaining species. An ART ANOVA allows for non-parametric testing for effects while still using regular ANOVA techniques (Durner 2019). This allows for a robust two-level analysis on non-normal ecological data (Guedes and Araujo et al. 2022; Khen et al. 2023). The same modelling approach was used for fish feeding mode functional groups (FMFGs) and total harvestable fish abundance. All data was analysed in R version 4.5.0.

## Results

A total of 24 species were identified across the two habitats and times in the 13 sites sampled. Of these, 18 were identified to species level and six to family level (Table 1). All taxa were grouped into trophic guilds using the estuarine feeding mode functional group (FMFG) classification by Elliott et al (2007), including zoobenthivores (ZB), piscivores (PV), detritivores (DV), herbivores (HV), omnivores (OV), and zooplanktivores (ZV). Fish were also grouped into harvested species as the sum of MaxN values for species harvested commercially in southeast Queensland (Gilby et al. 2021).

**Table 1.** Fish taxa observed across habitats and times, their FMFG, and the percentage of deployments in which they were present.

| Species / Taxon | Common Name | FM FG | Harvestable Species | Oyster / Day | Oyster / Night | Unstructured / Day | Unstructured / Night | Deployments present (%) |
| --- | --- | --- | --- | --- | --- | --- | --- | --- |
| *Acanthopagrus australis* | Yellowfin Bream | ZB | Y | Y | Y | Y | Y | 38.5 |
| *Ambassis marianus* | Estuary Glassfish | ZP |  | Y | Y | Y | Y | 32.7 |
| *Gerres oyena* | Common Silverbiddy | ZB | Y | Y |  | Y | Y | 9.6 |
| *Atherinomorus lacunosus* | Hardyhead | ZB |  | Y |  | Y | Y | 9.6 |
| *Scatophagus argus* | Spotted Scat | OV |  | Y | Y | Y |  | 7.7 |
| *Sillago ciliata* | Sand Whiting | ZB | Y |  | Y | Y | Y | 5.8 |
| *Mugil cephalus* | Sea Mullet | DV | Y | Y |  | Y |  | 5.8 |
| Myliobatidae | Rays | ZB |  | Y | Y |  |  | 5.8 |
| *Microcanthus strigatus* | East Australian Stripey | OV |  | Y |  |  |  | 5.8 |
| *Pelates sexlineatus* | Striped Grunter | HV |  | Y |  | Y |  | 3.8 |
| *Lutjanus argentimaculatus* | Mangrove Jack | PV | Y | Y |  | Y |  | 3.8 |
| Carangidae | Trevallies | PV | Y | Y |  |  |  | 3.8 |
| *Lutjanus russellii* | Moses Perch | PV | Y | Y |  |  |  | 3.8 |
| *Monodactylus argenteus* | Diamondfish | OV |  | Y |  |  | Y | 3.8 |
| Monocanthidae - unknown | Leatherjackets | OV | Y | Y |  |  |  | 3.8 |
| *Herklotsichthys castelnaui* | Southern Herring | ZP | Y |  |  | Y | Y | 3.8 |
| Diodontidae | Porcupinefishes | ZB |  | Y |  |  |  | 1.9 |
| *Epinephelus malabaricus* | Malabar Grouper | ZB | Y | Y |  |  |  | 1.9 |
| *Sphyraena obtusata* | Obtuse Barracuda | PV |  |  | Y |  |  | 1.9 |
| *Monocanthus chinensis* | Fantail Leatherjacket | OV | Y |  |  | Y |  | 1.9 |
| *Plotosus lineatus* | Eel-tailed Catfish | ZB |  |  | Y |  |  | 1.9 |
| Acanthuridae | Surgeonfishes | OV |  | Y |  |  |  | 1.9 |
| *Parachaetodon ocellatus* | Sixspine Butterflyfish | ZB |  | Y |  |  |  | 1.9 |
| Siganidae | Rabbitfishes | HV | Y | Y |  |  |  | 1.9 |
| **TOTAL taxa** |  |  | 12 / 24 | 19 / 24 | 7 / 24 | 11 / 24 | 7 / 24 |  |

Most species were observed at daytime, with 22 of the 24 recorded during daytime deployments (Table 1). Two species were observed only at night on oyster reefs - obtuse barracuda (*Sphyraena obtusata*) and eel-tailed catfish (*Plotusus lineatus*) - while 13 species were observed only during the day (Table 1). Twelve species were observed only at oyster reefs, in addition to 10 that were observed as users of both oyster and unstructured habitats (Table 1). Two species (*Monocanthus chinensis* and *Herklotsichthys castelnaui*) were recorded only on unstructured deployments. The most common species across all videos were yellowfin bream (*Acanthopagrus australis*; 38.5% of videos) and estuary glassfish (*Ambassis marianus*; 32.7%).

### Comparison of Fish Assemblages

Fish assemblages differed between day and night (PERMANOVA: F_time_ = 3.2951, p = 0.006) with the ordination plot for time suggesting that nighttime fish assemblages are a subset of those found during the day (Figure 4). There were no significant differences between oyster reef and unstructured habitats or the interaction (Table 2), with high overlap in the habitat NMDS plot suggesting little differentiation in fish assemblages (Figure 4).

**Figure 4.**
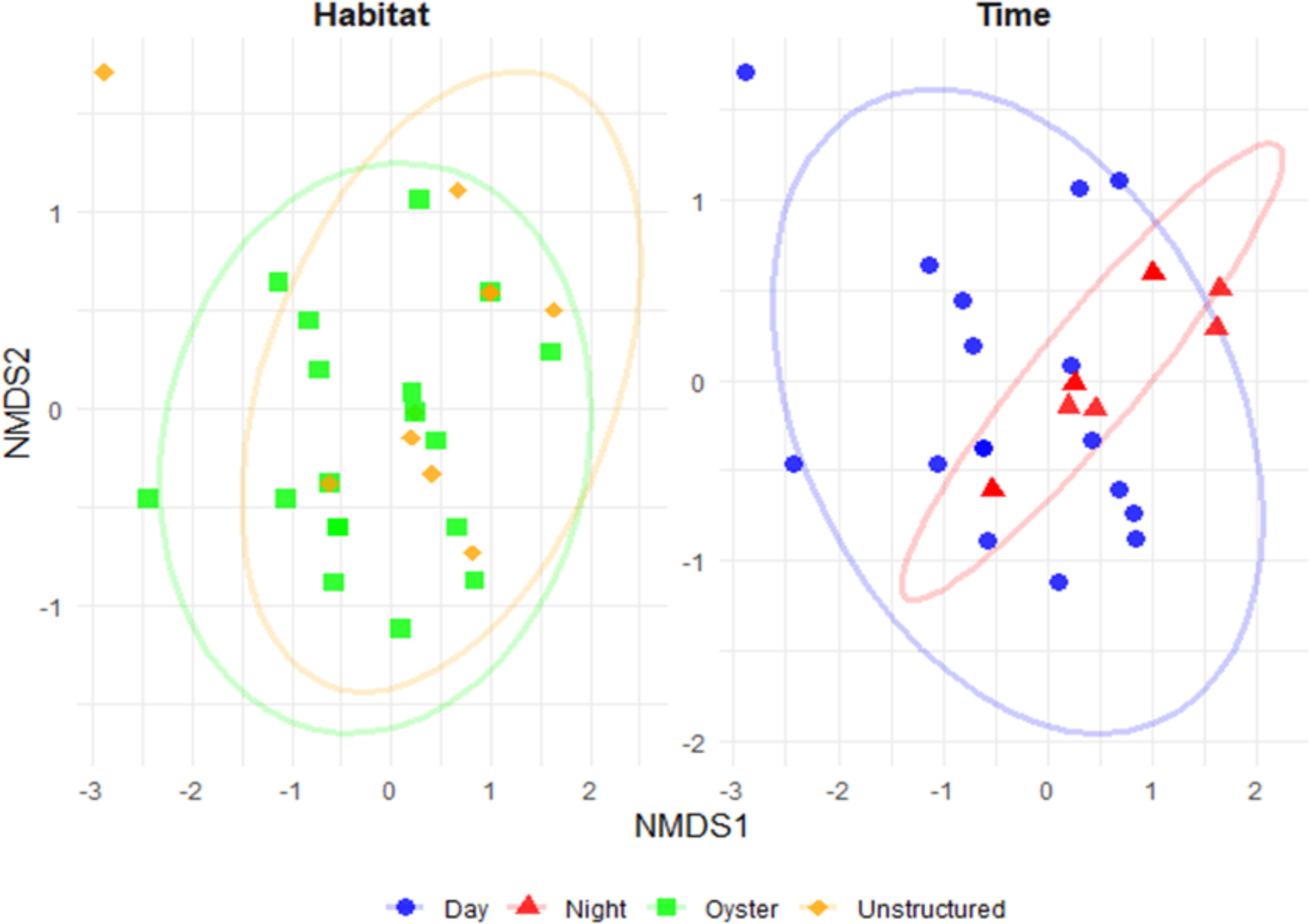
Habitat and time NMDS ordination plots of fish communities. Fish assemblages differed between day and night (F_time_ = 3.2951, p = 0.006). The assemblages did not differ between oyster and unstructured habitats.

**Table 2.** PERMANOVA results on time and habitat (Jaccard distance, stress = 0.076). There were eight species that occurred in only one deployment and were excluded from analysis. Boldface factors indicate statistical significance at p < 0.05.

|  | DF | Sum of Squares | F value | P-value |
| --- | --- | --- | --- | --- |
| Time | 1 | 0.9950 | 3.2951 | 0.006 |
| Habitat | 1 | 0.3514 | 1.1636 | 0.263 |
| Time:Habitat | 1 | 0.1179 | 0.3905 | 0.955 |
| Residual | 26 | 7.8509 |  |  |
| Total | 29 | 9.3151 |  |  |

Species richness and Shannon diversity showed similar trends: a significant interaction between time and habitat (Table 3), with the highest values at oyster reefs during the day (Figure 5; Table S2).

**Figure 5.**
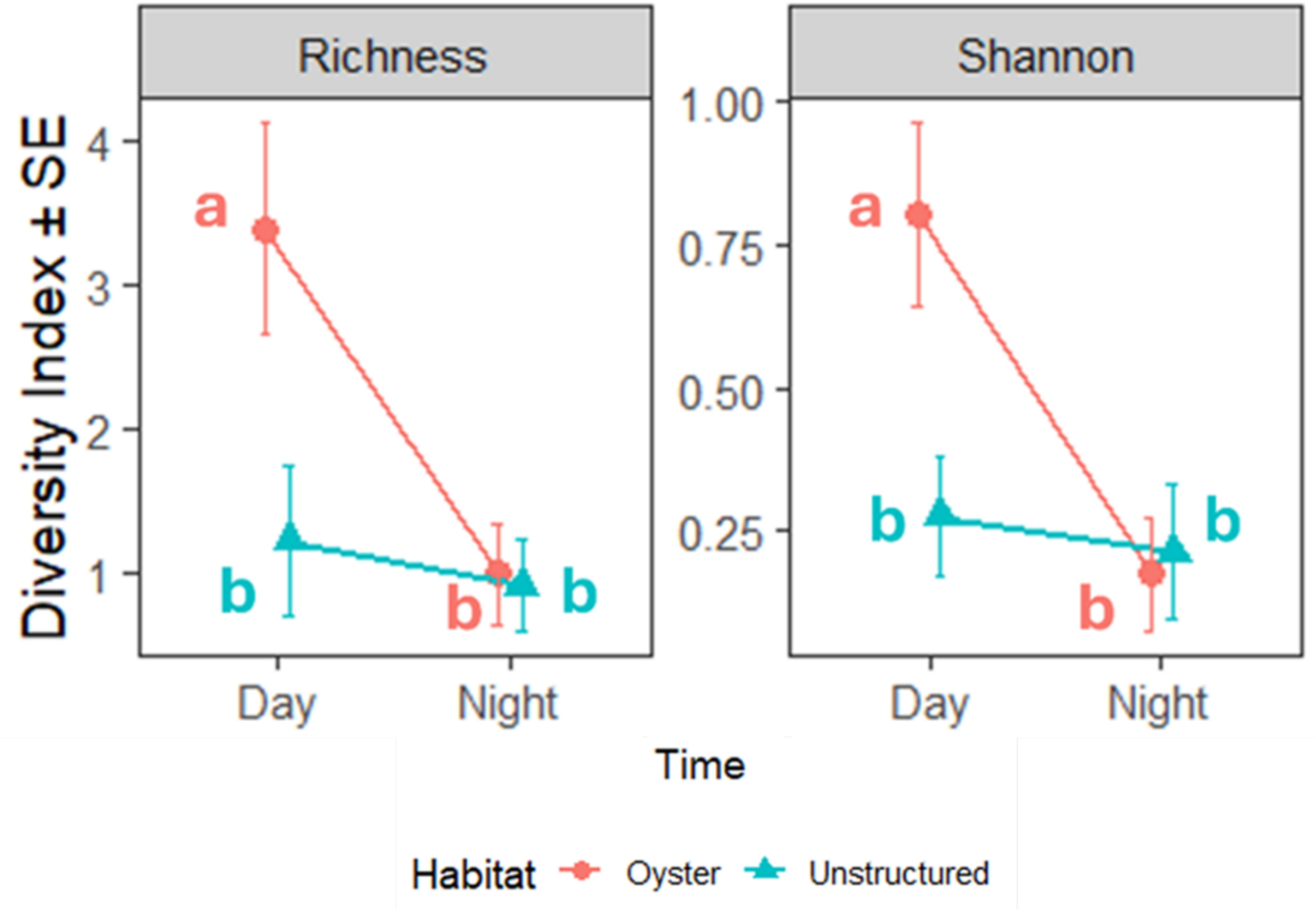
Shannon’s diversity and species richness of the whole fish community found on oyster reefs (red) and in unstructured sites (blue) for day and night deployments. Species richness and Shannon’s diversity showed a significant interaction effect between habitat and time.

**Table 3.** ANOVA table on linear mixed models for diversity indices, and species richness. Boldface factors indicate statistical significance at p < 0.05.

| *Species Richness (log-transformed)* |  |  |  |  |  |  |
| --- | --- | --- | --- | --- | --- | --- |
| **Time** | **2.1585** | **2.1585** | **1** | **36** | **8.5349** | **p < 0.01** |
| **Habitat** | **2.1190** | **2.1190** | **1** | **36** | **8.3784** | **p < 0.01** |
| **Time:Habitat** | **1.9426** | **1.9426** | **1** | **36** | **7.6811** | **p < 0.01** |
| *Shannon's Diversity Index* |  |  |  |  |  |  |
|  | Sum of Squares | Mean Squares | NumD F | DenDF | F-value | P-value |
| **Time** | **1.55398** | **1.55398** | **1** | **36** | **9.5737** | **p < 0.01** |
| **Habitat** | **0.78043** | **0.78043** | **1** | **36** | **4.8080** | **0.035** |
| **Time:Habitat** | **1.04904** | **1.04904** | **1** | **36** | **6.4629** | **0.015** |

### Species Abundances

We found significant interaction effects for the abundance of yellowfin bream (*A. australis*) and the common silverbiddy (*G. oyena*; Table S3). For both species, abundances were higher during the day in oyster reefs before converging at night with the lower abundances characteristic of unstructured habitats during both times (Figure 6). This pattern of high daytime abundance dropping at night - especially in oyster reefs - remained mostly consistent across most of the top nine most abundant taxa, although low abundances limited statistical inference. The two species that did not follow this trend were estuary glassfish (*A. marianus*) and eel-tailed catfish (*P. lineatus*; Figure 6). Unlike most other common species observed in this study, estuary glassfish showed similar abundances and trends across both times and habitats. Furthermore, eel-tailed catfish were only observed in one deployment and was not included in the PERMANOVA as its observed diel pattern may have been an outlier.

**Figure 6.**
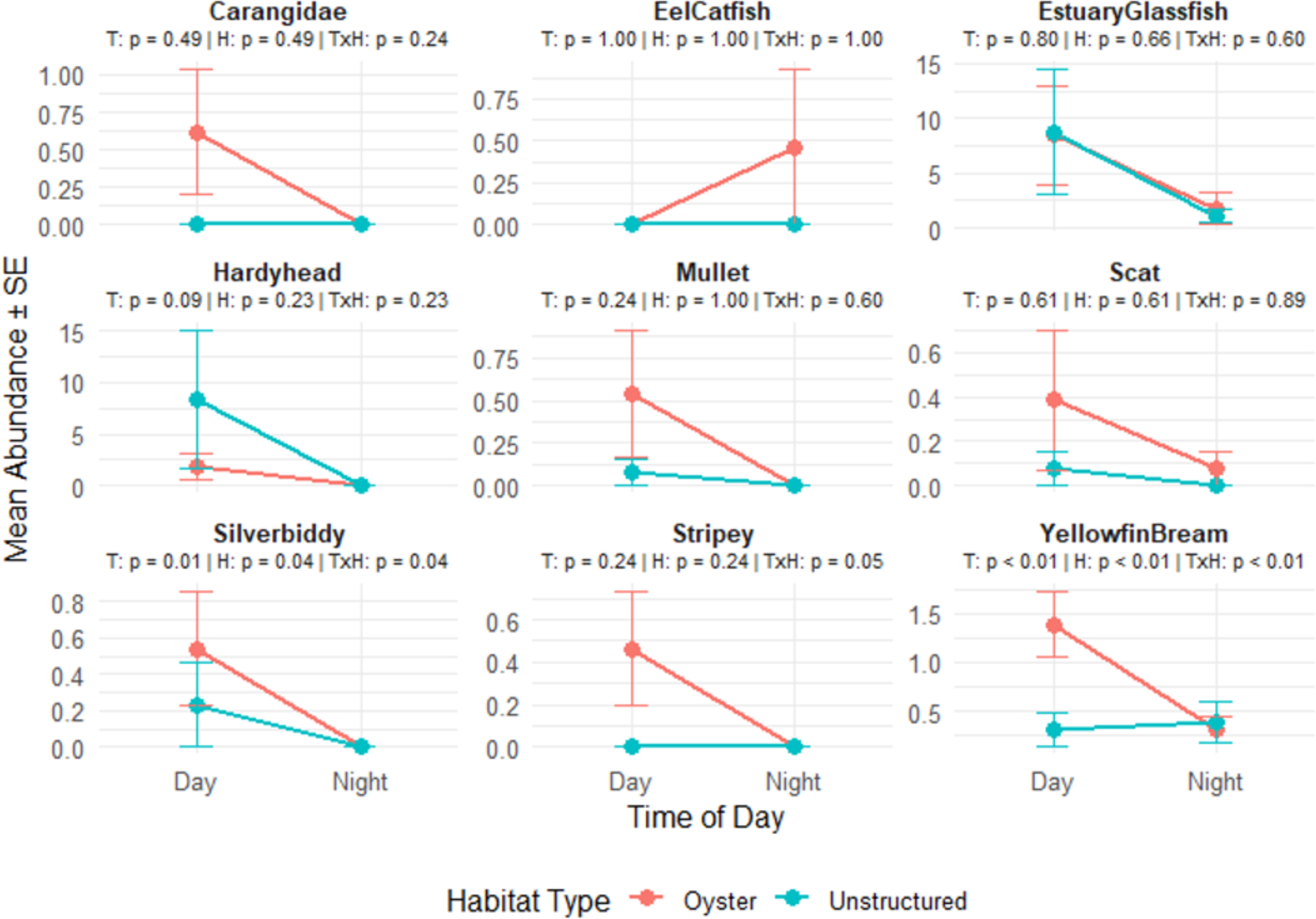
Mean abundance (MaxN per site +-SE) of the top 9 most prevalent species found on oyster reefs (red) and in unstructured sites (blue) for day and night deployments. P-values are provided, where T = time, H = habitat, and TxH = interaction.

### Functional Group and Harvestable Species Abundances

Omnivores exhibited a significant interaction effect (F_int_ = 9.3, p < 0.01), being most abundant at oyster reefs during the day (Figure 7; Table S6). Piscivores were more abundant at oyster reefs (F = 5.1, p = 0.03) and at night (F = 4.6, p = 0.04; Figure 7). Other groups showed no statistically significant differences.

**Figure 7.**
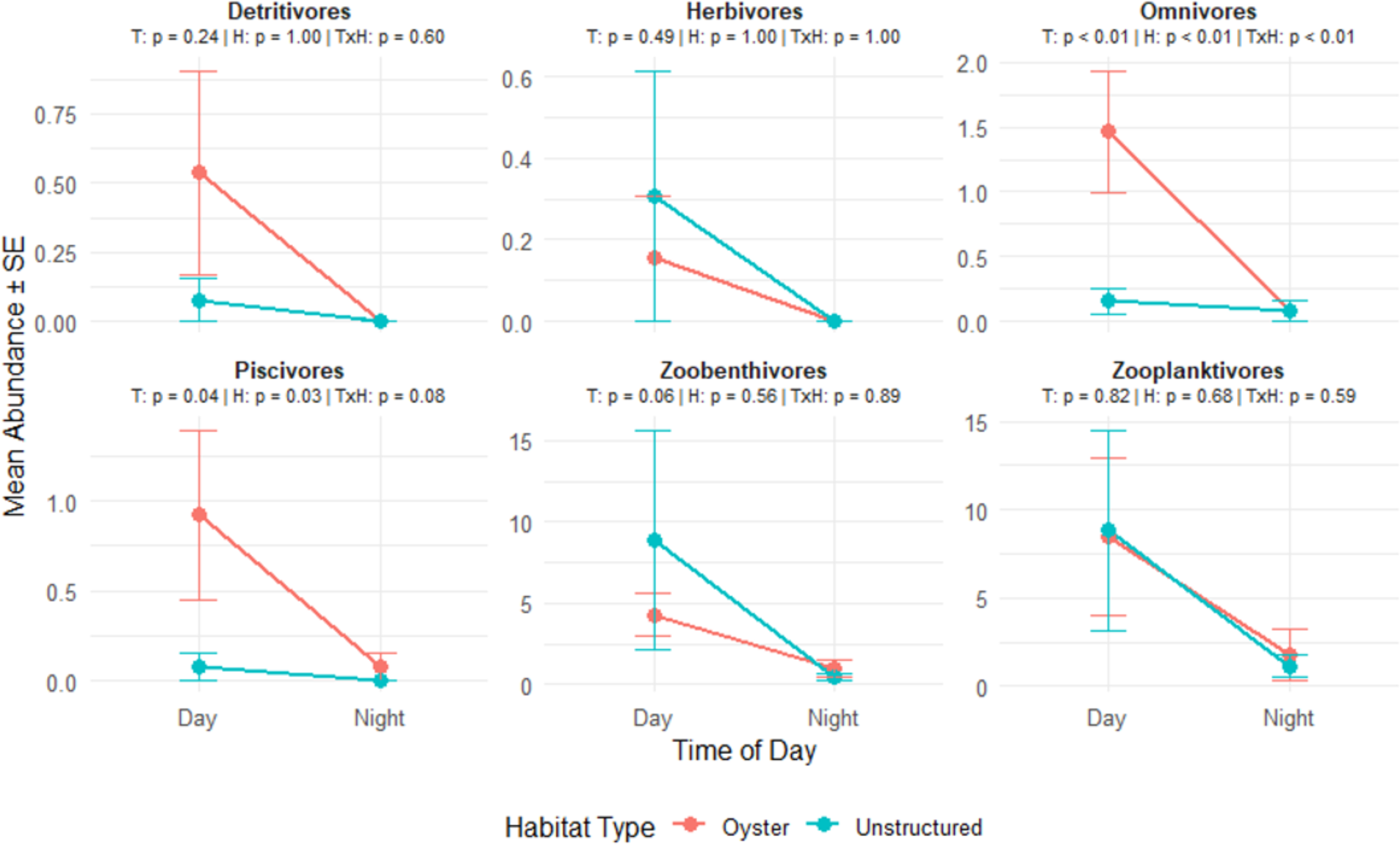
Mean abundance (MaxN per site; +-SE) of feeding mode functional groups found on oyster reefs (red) and in unstructured sites (blue) for day and night deployments. P-values are provided, were T = time, H = habitat, and TxH = interaction.

There was a significant interaction effect for harvestable fish species (F_int_ = 14.543, p < 0.01), with higher abundances during the day in oyster reefs, before converging at night with the low abundances in unstructured habitats at night (Figure 8; Table S8).

**Figure 8.**
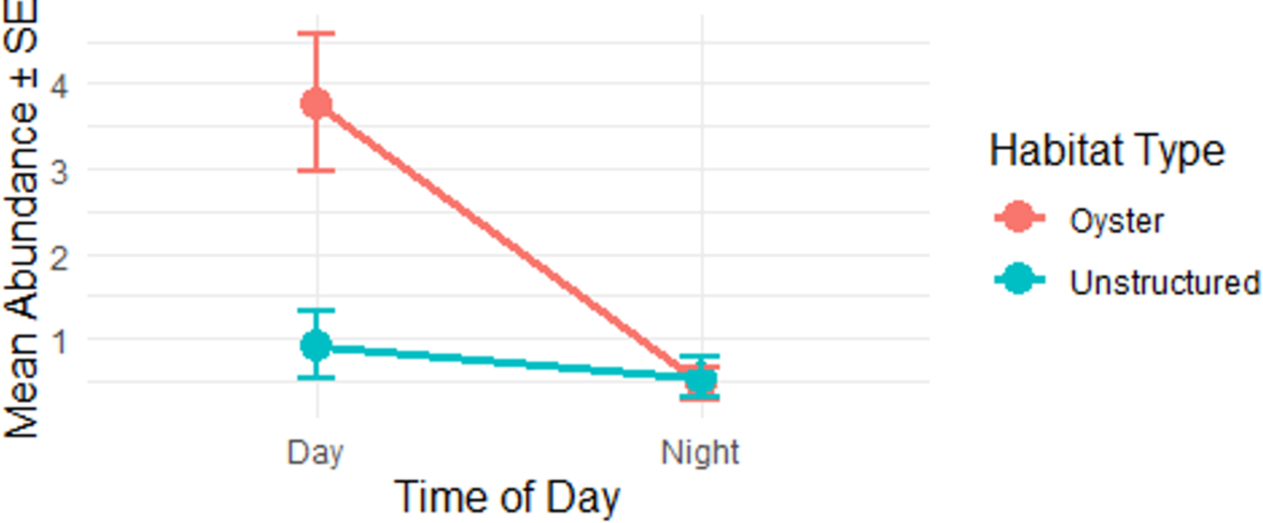
Mean abundance (MaxN per site; +-SE) of harvestable species found on oyster reefs (red) and in unstructured sites (blue) for day and night deployments.

## Discussion

### General Observations and Trends

Our data verifies previous knowledge that species abundances vary between day and night as fish engage in various behaviors and movements, leading to our observed changes in habitat use at the temporal level, where oyster reefs and adjacent unstructured adjacent habitats converged at night with regards to abundances and diversity (Magneville et al. 2022).

Unstructured habitats generally have lower species diversity and abundances during the day, likely due to reduced structure creating less niches, refuge, and opportunities for fish. Shannon diversity was highest in oyster reefs during the day. This is consistent with previous literature showing structurally complex oyster reefs supporting distinct communities and serving as habitats for economically important species (Pinnell et al. 2021; Martínez-Baena et al. 2022). While oyster reefs had higher species richness, diversity, and abundances during the day, we found two species, obtuse barracuda (*S. obtusata*) and eel-tailed catfish (*P. lineatus*) on reefs at night that were not present during the day. Fish abundances and diversity in both oyster reefs and unstructured habitats were generally low at night, suggesting that many fish species utilise alternative habitats throughout the diel cycle.

### Oyster Reefs are Key Fish Habitats

Our results are consistent with previous knowledge of oyster reefs playing an important role in habitat provision for fish in the estuarine seascape (Quan et al. 2011; Martínez-Baena et al. 2022). A total of 24 taxa were recorded in this study, of which 18 were identified to the species level. The majority of these (22 of 24) were found in oyster reefs, with 12 being exclusively recorded in the oyster habitats. Oyster reefs during the day were characterised by the highest species richness and diversity. This can be explained by their structural complexity, which may create a higher amount and variety of niches that different species can occupy within the reef (Pinnell et al. 2021). For example, our data showed that piscivores and omnivores were significantly higher in oyster reefs compared to unstructured habitats. Rays (family Myliobatidae), which can be indicators of a diverse fish assemblage and healthy ecosystems (Gilby et al. 2017), were only observed in oyster reefs during this study. These results underline the importance of oyster reefs as a key fish habitat, with a wider variety and rarer fish species more likely to be found in these areas.

### Oyster Reefs Provide for an Extensive Food Web

Oyster reefs can be productivity hotspots through the creation of organic matter and benthic microalgae (Quan et al. 2011; Engel et al. 2017), which serves as a food source for secondary consumers. Interestingly, we did not observe significantly higher abundances of herbivores or zoobenthivores inside the reef to support this. However, there were more individual zoobenthivore species that were recorded exclusively on the oyster reef. This may be the result of competition for resources inside the reef or the temporary attraction of highly mobile species. The productivity of oyster reefs also attracted higher abundances of omnivores, which may also increase competition for food. Interestingly, piscivores were significantly more abundant on oyster reefs, with many piscivorous species only observed in this habitat. These included the Moses Perch (*Lutjanus russellii*), Obtuse Barracuda (*Sphyraena obtusata*) and trevallies (family Carangidae). Predation probability in estuaries can be shaped by either predator species richness or predator abundance (Mosman et al. 2023), and both of these were observed in oyster reefs during this study. This suggests that predation is a key process occurring in these habitats, certainly during the day and likely across multiple trophic levels. With higher piscivore populations on oyster reefs, it is likely that predation also serves as a top-down control in these habitats. The presence of these higher-level predators may suggest that oyster reefs provide for a diverse food web that extends beyond the benthic matter and shellfish that composes the reef structure.

### The Night Brings Different Trends

Time played a significant role in structuring fish assemblages, including on species richness and diversity, individual species abundances, and feeding mode functional groups. In contrast to previous studies on other habitats that have found higher fish abundances and/or biomasses at night in other structured habitats (Griffiths 2001; Guest et al 2003; Hagan and Able 2007; Matheson Jr. et al. 2017; McSpadden et al. 2023), we found that fish abundances and diversity were lower at night in oyster reefs. Previous studies have also suggested that some species might be documented at night (Griffiths 2001; Hagan and Able 2007), and indeed we found two species that were documented exclusively at night - the eel-tailed catfish (*P. lineatus*) and obtuse barracuda (*S. obtusata*). This emphasises the role that night sampling plays in documenting a more comprehensive and wider range of habitat users.

Nighttime seems to “equalise” the two habitats, as abundance and diversity metrics converged at night after being different during the day. This is consistent with our initial prediction of an interaction between habitat and time in structuring species assemblages in oyster reefs. This may be due to a reduction of overall fish abundance and the number of species present in the entire intertidal zone due to reduced activity at night. Fish may be resting or retreating to deeper waters at night (Taylor et al. 2013), outside of the intertidal habitats surveyed. If there are fewer fish in the intertidal zone, then the two habitats we measured would be more similar, thus explaining their convergence at nighttime.

Although there is a general decline of abundance and diversity at night in oyster reefs, the magnitude and pattern of this reduction is not the same across all species or trophic guilds, suggesting differing diel uses of oyster reefs by different trophic guilds. Furthermore, while we did not observe a higher relative abundance of piscivores at night, it is possible that fish modify behavior in response to increased predation risk at night (Taylor et al. 2013; Bosiger and McCormick 2014), thus resulting in lower abundances. These results can provide insights to temporal niche partitioning, where animals exploit different times of day to reduce risk of predation and competition for food (Kronfeld-Schor and Dayan 2003). Larger and more dominant species are likely to forage at times that are best suited to their physiology, while other less dominant species may be forced to feed at other times to avoid them (Lear et al. 2021). Our results suggest that although oyster reefs generally harbor more diversity, there is still consistency across the entire coastal seascape in the pattern of the dominance of only a few - and likely well-adapted - species at night.

The abundance of piscivores was significantly lower at night. This result is supported by our observations of Moses Perch (*L. russellii*) and Mangrove Jack (*L. argentimaculatus*), which were exclusively recorded during daytime deployments despite knowledge of their behaviour as a nocturnal predator (Laprise and Blaber 1992; Zagars et al. 2012). This is surprising given that many higher order predators tend to increase in abundance at night in other habitats (Griffiths 2001; Hagan and Able 2008; Castillo-Rivera et al. 2010). However, it is possible that these fish were still active at night but switched behaviour to more furtive hunting and ambush strategies that may have made them harder to detect. Another possibility is that oyster reefs do not serve as feeding grounds during the night and that piscivores move to other areas to feed during this period. A significant interaction between habitat and time was observed for omnivores, with lower abundances at night in oyster reefs but not in unstructured habitats.

Observations of omnivores in other aquatic habitats have found that their foraging behaviour can change at a diel scale (Kwak et al. 2006; Nakagawa et al. 2012; Nakagawa 2023). As omnivores have a wider diet range, it is possible that their food preference switches at a diel pattern. This is consistent with the foraging arena theory that proposes the idea of animals having to balance predation risk with food reward (Ahrens et al. 2011), and with these values altered at night, fish may switch behaviors to exploit areas that they would not normally frequent during the day or avoid them altogether.

Darkness may alter the value of structure to fish. While it can be a source of refuge and protection, structural complexity can also provide ambush areas that prey may want to avoid (Michel et al. 2020). This risk of ambush combined with the higher difference in physiological adaptations between predators and prey at night may cause fish to respond by either reducing their activity or switching habitats during the night. Birds, which are key predators in shallow coastal areas (Gawlik 2002; Steinmetz et al. 2003; Žydelis and Kontautas 2008), may be less active at dark and a possible lowered risk of avian predation at night may further reduce the protection value of structure. This research is unique in that it sampled intertidal habitats where fish are given the choice of recolonising oyster reefs or unstructured habitats at a diel pattern. Unlike subtidal zones where fish can choose to stay in a given habitat for an indefinite period of time, intertidal areas are subject to more pressure and most fish are forced out of these habitats with the tidal cycle. The benefits and risks associated with oyster reefs may change throughout the day, with the structure of oyster reefs possibly creating a riskier habitat for most fish species at night. As fish must balance the risk of predation with other behaviors such as foraging (Ahrens et al. 2011), their decision of which habitat to utilise at night may change. When the tide returns under the cover of darkness, fish may adapt their behavior to reflect these decisions and maximise their chances of survival. This may mean avoiding areas that would otherwise be more beneficial during the daytime or staying in deeper waters and avoiding the intertidal zone altogether.

### Observations in Individual and Harvestable Fish Species

Across all deployments, yellowfin bream (*A. australis*) was the most commonly occurring species (38.5% of all deployments) and was observed in all four habitat-time combinations. Highly abundant species can reflect habitat use (McAneney et al. 2025) and yellowfin bream exhibit high side fidelity (Gannon et al. 2015), which suggests that our observations here are indicative of oyster reefs playing a role in its life history. This is not surprising, as bream is a ubiquitous and highly adaptable fish (Taylor et al. 2018) with a wide diet range that can be found across the estuarine seascape (Gannon et al. 2015). Their increased abundance during the daytime coincides with higher bream activity previously recorded in intertidal mangroves during the day, where shallower waters likely provide protection from predators (Payne et al. 2013; Gannon et al. 2015). During the night, bream may then migrate to deeper waters to hide from predators (Taylor et al. 2013), explaining the reduction in observed abundance during our night surveys. It is possible that this diel switch of behavior also holds true for other species.

The common silverbiddy (*G. oyena*) was the only other species to display a significant interaction effect. Although they can accommodate a wide range of food items, common silverbiddy can experience an ontogenetic shift in food items towards crustaceans and zoobenthic sources (Putri et al. 2022) which would likely be found in high abundances in oyster reefs. This may suggest that the individuals observed on the oyster reef may have been older. Also worth noting is the marginally significant interaction effect (p = 0.0518) observed for the East Australian stripey (*M. strigatus*). This genus is a reef associated taxon (Tea and Gill 2020) that can clean parasites off reef fish, including the yellowfin bream and moses perch observed in our surveys (Ebner 2025). This would explain the similar diel and habitat patterns to the aforementioned host species, which were also observed in this study.

Harvestable species as a group were more abundant in oyster reefs during the daytime. affirming their role as fishery hotspots as suggested in previous literature (Grabowski et al. 2012; Gilby et al. 2021). Although this study was not able to obtain biomass estimates, it presents local evidence that can guide protection and restoration efforts (Gaylard et al. 2020) by providing a more comprehensive list of fish species using intertidal oyster reefs and information on diel differences in their habitat use. This also provides further reason to pursue economic valuations of oyster reefs to improve policy (Barbier 2017). Obtaining biomass and monetary estimations of the fishery value of oyster reefs will likely become easier in the future as cheaper and more feasible methods are developed, including easier methods to size fish to subsequently estimate biomass from known length-weight relationships.

### Potential Future Applications of Monocular Estimation Technology

The data that could be analysed by monocular estimation for depth and sizes was too limited to allow for any meaningful comparisons. Nonetheless, this study shows promise in the future use of monocular estimating technology. These units are smaller and easier to deploy than traditional stereo cameras, and do not require expensive licenses to process data. Although our ability to make meaningful size comparisons was limited by a small dataset, environmental conditions, and inadequacies with initial set-ups, it is reasonable to assume that the artificial intelligence model will improve as more and better data is used to train it. For instance, we were able to improve calibration and field techniques here to achieve more reliable estimates and we anticipate that these changes will aid in future studies. As numerous commercially important species were observed in our videos, this technology will enable us to quantify sizes to estimate biomass and model fisheries production. This would be useful to determine the economic value and contribution of oyster reefs to fishery biomass, as well as the specific role it plays in the life cycles of important species.

### Implications for Conservation and Future Directions

We observed several important species in our surveys of oyster reefs, including a high relative abundance of the economically important yellowfin bream and common silverbiddy (Momtaz and Gladstone 2008; El Ganainy et al. 2020; Putri et al. 2022). While it is accepted that species need to be protected throughout their entire life history, this study also emphasises the importance of the protection of habitats used by species at a diel scale. Our results show that previously accepted knowledge of oyster reefs as fish habitat is not as simple as initially expected. As nighttime brings different patterns and lowered abundances on the reef, it is likely that fish migrate to adjacent habitats, including nearby deeper waters, for shelter. We currently know that habitat connectivity and seascape context can shape fish assemblages in coastal ecosystems, including oyster reefs (Whitfield 2017; Gilby et al. 2018; Gilby et al. 2019; Jones et al. 2021). However, this relationship between habitats is less clear at night; we have found that abundances are reduced at night, but further investigation is required to understand where the fish migrate during this period. Protection of oyster reefs alone may not be enough - buffer zones of surrounding and deeper waters may be necessary to cover the range that a fish might use on a day-to-day basis. From a research perspective, we have found that while nighttime surveys may not necessarily show higher abundances of species, they still identify unique species and reflect important shifts in how key inhabitants use ecosystems.

### Conclusion

To our knowledge, this study provides the first comparison of day and night surveys of fish assemblages in intertidal oyster reefs using non-invasive methods. During the day, we observed higher abundances of piscivore and omnivore feeding guilds using the oyster reef alongside yellowfin bream and common silverbiddy, and the overall abundance of harvestable species. Patterns during the day do not match those at night, with lowered diversity and overall abundance observed during the nighttime. Furthermore, this reduction is more pronounced in oyster reefs than in unstructured habitats. This interaction between habitat and time of day suggests that fish use of intertidal oyster reefs changes more dramatically over the diel cycle. Further insights can be gained as monocular depth estimation and sizing technology improves, with biomass estimations and fish life stage estimations likely to be possible in the near future. Our results emphasise the consideration of nighttime interactions and shifts in the study and management of coastal ecosystems.

## Acknowledgments

MS was supported by a funded by Australian Research Council (ARC) Discovery Early Career Researcher Award no. DE220100079. Shakya Fernando assisted in the data processing and use of monocular estimation technology. This research was supported by use of the Nectar Research Cloud and by FishID.

## Author Contributions

DRG, MS, CH, RMC conceptualised the study; Field work was conducted by DRG, MS and resourced by MS, RMC; Videos were processed by DRG and CH; All authors interpreted results; analysed the data; Manuscript writing was led by DRG with contributions from all authors. All authors approve submission of this manuscript.

## Competing Interest Statement

The authors disclose no competing interests.

## Notes

### Competing Interest Statement

The authors have declared no competing interest.

