## Supplementary Material for "Shedding light on day-night habitat use of oyster reefs by fish"

### SUPPLEMENTARY MATERIALS

Supplementary Table S1. Coordinates of oyster reef sample sites in southeast Queensland.

| Site Number | Coordinates |
|---|---|
| 1 | -27.042250°, 153.115930° |
| 2 | -27.047671°, 153.121135° |
| 3 | -27.051139°, 153.120120° |
| 4 | -27.054033°, 153.123752° |
| 5 | -27.085840°, 153.135578° |
| 6 | -27.085936°, 153.136591° |
| 7 | -27.086110°, 153.133878° |
| 8 | -27.086828°, 153.136006° |
| 9 | -27.876444°, 153.383983° |
| 10 | -27.877727°, 153.382930° |
| 11 | -28.098961°, 153.456896° |
| 12 | -28.106855°, 153.450117° |
| 13 | -28.108321°, 153.450358° |

Supplementary Table S2. Post-hoc pairwise comparisons of Shannon diversity index and species richness.

| **Pairwise Comparison** | **Estimate** | **SE** | **DF** | **T. Ratio** | **P-value** |
|---|---|---|---|---|---|
| *Shannon's Diversity* | | | | | |
| Oyster Day / Oyster Night | 0.6298 | 0.158 | 36 | 3.986 | 0.0017 |
| Oyster Day / Unstructured Day | 0.5291 | 0.158 | 36 | 3.348 | 0.0099 |
| Oyster Day / Unstructured Night | 0.5908 | 0.158 | 36 | 3.738 | 0.0034 |
| Oyster Night / Unstructured Day | - 0.1007 | 0.158 | 36 | - 0.637 | 0.9192 |
| Oyster Night / Unstructured Night | - 0.0391 | 0.158 | 36 | - 0.247 | 0.9946 |
| Unstructured Day / Unstructured Night | 0.0617 | 0.158 | 36 | 0.390 | 0.9795 |
| *Species Richness (log-transformed)* | | | | | |
| Oyster Day / Oyster Night | 0.79405 | 0.197 | 36 | 4.026 | 0.0015 |
| Oyster Day / Unstructured Day | 0.79029 | 0.197 | 36 | 4.006 | 0.0016 |
| Oyster Day / Unstructured Night | 0.81121 | 0.197 | 36 | 4.113 | 0.0012 |
| Oyster Night / Unstructured Day | - 0.00375 | 0.197 | 36 | - 0.019 | 1.0000 |
| Oyster Night / Unstructured Night | 0.01716 | 0.197 | 36 | 0.087 | 0.9998 |
| Unstructured Day / Unstructured Night | 0.02092 | 0.197 | 36 | 0.106 | 0.9996 |

Supplementary Table S3. Statistical Results for ART ANOVAs on Abundant Species.

| | **F-statistic** | **DF** | **Residual DF** | **P-value** |
|---|---|---|---|---|
| **Yellowfin Bream (*A. australis*), n = 31** | | | | |
| Time | 9.7058 | 1 | 36 | 0.004 |
| Habitat | 12.3243 | 1 | 36 | 0.001 |
| Time:Habitat | 10.5174 | 1 | 36 | 0.003 |
| **Common Silverbiddy (*G. oyena*), n = 10** | | | | |
| Time | 7.3356 | 1 | 36 | 0.01 |
| Habitat | 4.7654 | 1 | 36 | 0.04 |
| Time:Habitat | 4.7654 | 1 | 36 | 0.04 |
| **Hardyhead (*Atherinomorus lacunosus*), n = 133** | | | | |
| Time | 3.1324 | 1 | 36 | 0.09 |
| Habitat | 1.5235 | 1 | 36 | 0.23 |
| Time:Habitat | 1.5235 | 1 | 36 | 0.23 |
| **Estuary Glassfish (*A. marianus*), n = 261** | | | | |
| Time | 0.063106 | 1 | 36 | 0.80 |
| Habitat | 0.201531 | 1 | 36 | 0.66 |
| Time:Habitat | 0.274981 | 1 | 36 | 0.60 |

Supplementary Table S4. Statistical Results of Fisher's Test for Rare Species

| | **P-Value** |
|---|---|
| **Moses Perch (*L. russellii*), n = 3** | |
| Time | 0.4902 |
| Habitat | 0.4902 |
| Time:Habitat | 0.2353 |
| **Mangrove Jack (*L. argentimaculatus*), n = 2** | |
| Time | 0.4902 |
| Habitat | 1 |
| Time:Habitat | 1 |
| **Rabbitfish (family Siganidae), n = 1** | |
| Time | 1 |
| Habitat | 1 |
| Time:Habitat | 1 |
| **Trevallies (family Carangidae), n = 8** | |
| Time | 0.4902 |
| Habitat | 0.4902 |
| Time:Habitat | 0.2353 |
| **Rays (family Myliobatidae), n = 3** | |
| Time | 1 |
| Habitat | 0.2353 |
| Time:Habitat | 0.6024 |
| **Malabar Grouper (*E. malabricus*), n = 1** | |
| Time | 1 |
| Habitat | 1 |
| Time:Habitat | 1 |

| | |
|---|---|
| **Mullet (*M. cephalus*), n = 8** | |
| Time | 0.2353 |
| Habitat | 1 |
| Time:Habitat | 0.6024 |
| **Sand Whiting (*S. ciliata*), n = 4** | |
| Time | 1 |
| Habitat | 1 |
| Time:Habitat | 1 |
| **Porcupinefish (family Diodontidae), n = 1** | |
| Time | 1 |
| Habitat | 1 |
| Time:Habitat | 1 |
| **Stripey (*M. strigatus*), n = 6** | |
| Time | 0.2353 |
| Habitat | 0.2353 |
| Time:Habitat | 0.0518 |
| **Obtuse Barracuda (*S. obtusata*), n = 1** | |
| Time | 1 |
| Habitat | 1 |
| Time:Habitat | 1 |
| **Striped Grunter (*P. sexlineatus*), n = 5** | |
| Time | 0.4902 |
| Habitat | 1 |
| Time:Habitat | 1 |
| **Fantail Leatherjacket (*M. chinensis*), n = 1** | |

| | |
|---|---|
| Time | 1 |
| Habitat | 1 |
| Time:Habitat | 1 |
| **Leatherjackets (family Monocanthidae), n = 3** | |
| Time | 0.4902 |
| Habitat | 0.4902 |
| Time:Habitat | 0.2353 |
| **Spotted Scat (*S. argus*), n = 7** | |
| Time | 0.6098 |
| Habitat | 0.6098 |
| Time:Habitat | 0.8945 |
| **Eel-Tailed Catfish (*P. lineatus*), n = 6** | |
| Time | 1 |
| Habitat | 1 |
| Time:Habitat | 1 |
| **Diamondfish (*M. argenteus*), n = 5** | |
| Time | 1 |
| Habitat | 1 |
| Time:Habitat | 1 |
| **Surgeonfishes (family Acanthuridae), n = 1** | |
| Time | 1 |
| Habitat | 1 |
| Time:Habitat | 1 |
| **Sixspine Butterflyfish (*P. ocellatus*), n = 1** | |
| Time | 1 |

| | |
|---|---|
| Habitat | 1 |
| Time:Habitat | 1 |
| **Southern Herring (*Herklotsichthys castelnaui*), n = 2** | |
| Time | 1 |
| Habitat | 0.4902 |
| Time:Habitat | 1 |

Supplementary Figure S5. Interaction plots for remaining less abundant taxa. All taxa displayed in this plot were analysed using Fisher's test.

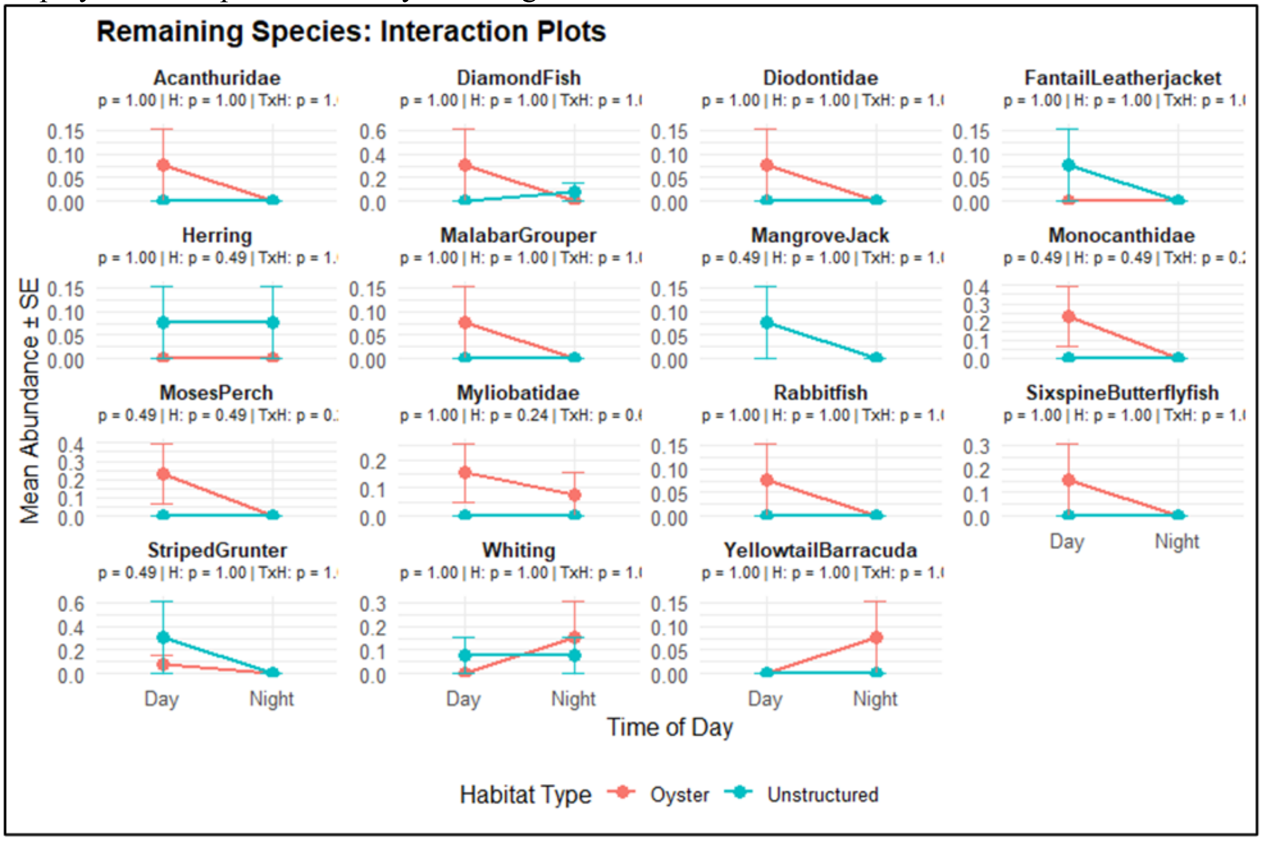

Supplementary table S6. Statistical results for ART ANOVA on abundant FMFGs.

| | F-statistic | DF | Residual DF | P-value |
|---|---|---|---|---|
| **Zoobenthivores, n = 191** | | | | |
| Time | 3.740094 | 1 | 36 | 0.061 |
| Habitat | 0.349713 | 1 | 36 | 0.558 |
| Time:Habitat | 0.020044 | 1 | 36 | 0.888 |
| **Piscivores, n = 14** | | | | |
| Time | 4.5887 | 1 | 36 | 0.039 |
| Habitat | 5.0642 | 1 | 36 | 0.031 |
| Time:Habitat | 3.2035 | 1 | 36 | 0.082 |
| **Omnivores, n = 23** | | | | |
| Time | 13.8230 | 1 | 36 | $p < 0.01$ |
| Habitat | 9.3005 | 1 | 36 | $p < 0.01$ |
| Time:Habitat | 9.3005 | 1 | 36 | $p < 0.01$ |
| **Zooplanktivores, n = 263** | | | | |
| Time | 0.050502 | 1 | 36 | 0.823 |
| Habitat | 0.170870 | 1 | 36 | 0.681 |
| Time:Habitat | 0.298534 | 1 | 36 | 0.588 |

Supplementary Table S7. Statistical Results for Fisher's Test on less abundant FMFGs.

| | |
|---|---|
| **Detritivores, n = 8** | |
| Time | 0.2353 |
| Habitat | 1 |
| Time:Habitat | 0.6024 |
| **Herbivores, n = 6** | |
| Time | 0.4902 |
| Habitat | 1 |
| Time:Habitat | 1 |

Supplementary table S8. Statistical results for ART ANOVA on harvestable species (n = 74)

| | **F-statistic** | **DF** | **Residual DF** | **P-value** |
|---|---|---|---|---|
| Time | 18.076 | 1 | 36 | 0.00014377 |
| Habitat | 10.226 | 1 | 36 | 0.00288473 |
| Time:Habitat | 14.543 | 1 | 36 | 0.00051755 |
